# Tumor genotype dictates radiosensitization after *Atm* deletion in brainstem gliomas

**DOI:** 10.1101/2020.08.24.262642

**Authors:** Katherine M. Deland, Bryce F. Starr, Joshua S. Mercer, Jovita Byemerwa, Donna M. Crabtree, Nerissa T. Williams, Lixia Luo, Yan Ma, Mark Chen, Oren J. Becher, David G. Kirsch

## Abstract

Diffuse intrinsic pontine glioma (DIPG) kills more children than any other type of brain tumor. Despite clinical trials testing many chemotherapeutic agents, radiotherapy remains the standard treatment. Here, we utilized Cre/loxP technology to show that deleting *Ataxia telangiectasia mutated* (*Atm*) in primary mouse models of DIPG can enhance tumor radiosensitivity. Genetic deletion of *Atm* improved survival of mice with *p53* deficient but not *p53* wild-type gliomas following radiotherapy. Similar to patients with DIPG, mice with *p53* wild-type tumors had improved survival after radiotherapy independent of *Atm* deletion. *p53* wild-type tumor cell lines induced proapoptotic genes after radiation and repressed the NRF2 target, *Nqo1*. Tumors lacking *p53* and *Ink4a/Arf* expressed the highest level of *Nqo1* and were most resistant to radiation, but deletion of *Atm* enhanced the radiation response. These results suggest that tumor genotype may determine whether inhibition of ATM during radiotherapy will be an effective clinical approach to treat DIPGs.

## Introduction

Diffuse intrinsic pontine glioma (DIPG), also referred to as high-grade brainstem glioma, is an incurable cancer that originates in the pons and occurs primarily in children. Focal radiation therapy is the current standard of care for children with DIPG, as the anatomic location of the tumor and its infiltrating growth patterns preclude surgery. Furthermore, no chemotherapeutic agents have improved outcome over radiotherapy alone. Although neurological symptoms typically improve following radiotherapy, the tumors invariably recur. The median survival for children with DIPG is less than one year and fewer than 10% of patients survive two years from diagnosis (1, 2). Therefore, DIPG is the major cause of brain cancer-related deaths in children (1). One approach to improving the survival of children with DIPG would be to develop effective strategies to overcome the radiation resistance of brainstem gliomas.

As damage to DNA is the primary cause of cell death by ionizing radiation (3), the proteins that sense DNA damage and signal the cell to repair damaged DNA play an essential role in maintaining genomic stability and regulating radiation response. Therefore, small molecule drugs that interfere with the DNA damage response represent a promising strategy for enhancing the ability of radiotherapy to kill cancer cells (4). Furthermore, a recent RNAi screen in patient-derived DIPG cells identified several kinases involved in the DNA-damage response whose inhibition selectively impaired survival following radiation exposure (5). One such kinase, Ataxia Telangiectasia Mutated (ATM) is recruited to sites of damage and activated through autophosphorylation (6–8). ATM phosphorylates numerous downstream effector proteins to orchestrate DNA repair (e.g. p53, MRE11, RAD50, BRCA1) (9) and contributes indirectly to DNA repair by phosphorylating proteins that facilitate cell cycle arrest and chromatin relaxation (e.g. CHK2, KAP1) (10–13).

Consistent with the role of ATM as a critical regulator of the DNA damage response, patients with ataxia telangiectasia that have two germline mutations in *ATM* are hypersensitive to radiation (14, 15). Similarly, *Atm^-/-^* mice die when exposed to sublethal radiation doses (16). Therefore, pharmacological inhibition of ATM kinase represents a rational approach to radiosensitize human tumor cells (17, 18). Importantly, the development of an ATM inhibitor that penetrates the blood-brain-barrier (19) has enabled a phase I clinical trial of adult patients with glioblastoma testing concurrent ATM inhibition and radiotherapy (NCT03423628). If this clinical trial shows that ATM inhibition during radiotherapy is safe, then testing this approach in DIPG patients could follow if preclinical studies demonstrate that targeting ATM improves survival in pre-clinical models of DIPG.

Here, we utilized primary genetically engineered mouse models (GEMMs) of brainstem glioma with different genotypes to explore the impact of tumor genotype and *Atm* deletion on tumor response to radiotherapy. We found that *p53* deficient but not *p53* wild-type brainstem gliomas (driven by deletion of *Ink4a/Arf)* were radiosensitized by disruption of ATM signaling. Consistent with data from human DIPG patients (5), mice with *p53* wild-type brainstem gliomas exhibited an enhanced radiation response. The inherent radiosensitivity of these *Ink4a/Arf* deficient tumors was mediated by p53 signaling and associated with repression of the NRF2 pathway gene *Nqo1.* We showed that deletion of *p53* in the *Ink4a/Arf* deficient gliomas resulted in overexpression of *NAD(P)H quinone dehydrogenase 1 (Nqo1)* and significantly increased radiation resistance *in vivo.* Importantly, deletion of *Atm* in the radioresistant brainstem gliomas lacking *p53* improved the survival of tumor-bearing mice treated with radiotherapy. Our findings suggest that ATM is a promising target for the radiosensitization of DIPGs and that the outcomes of clinical trials testing concurrent ATM inhibition and radiotherapy should be stratified according to *p53* mutational status.

## Results

### Generating primary brainstem gliomas

To investigate whether disruption of *Atm* can modulate the radiosensitivity of DIPGs, we employed *Cre-loxP* and RCAS-TVA technology to generate primary brainstem gliomas as previously described (20). Chicken fibroblast cells producing various RCAS retroviruses were injected into the brainstem of neonatal *Nestin^TVA^* mice, in which *Nestin-expressing* progenitor cells express the RCAS cognate tumor virus A (TVA) receptor (Figure 1A). We initiated glioma development by using RCAS-PDGFB to overexpress the *platelet-derived growth factor subunit B* (PDGFB) oncogene and RCAS-Cre to delete floxed alleles of various tumor suppressors (Figure 1B). In addition to promoting tumorigenesis, Cre recombinase also deleted either one or two floxed alleles of *Atm* in order to modulate the radiosensitivity of tumor parenchymal cells.

**Figure 1.**
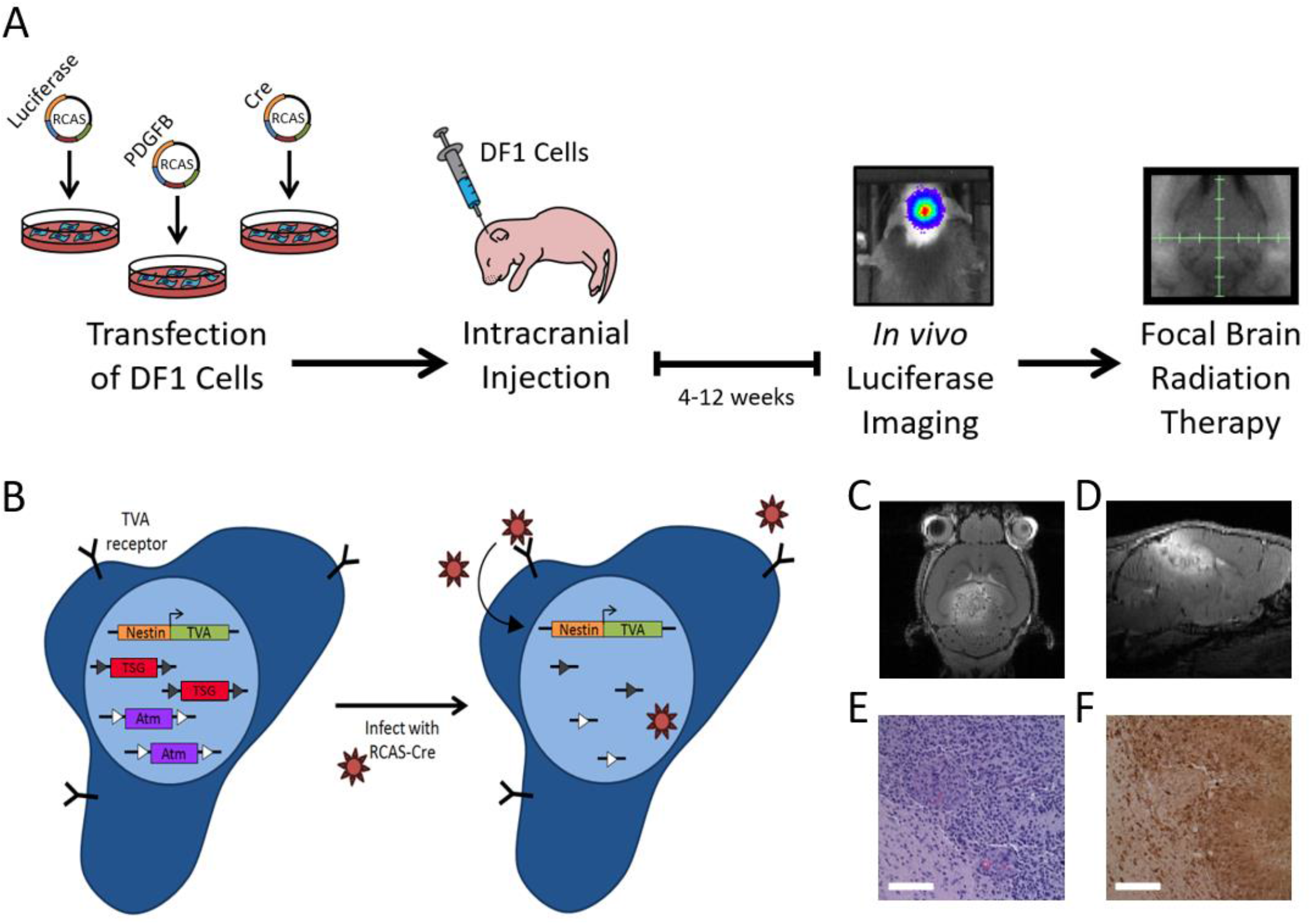
Genetically Engineered Mouse Model of Brainstem Glioma. (**A**) DF1 chicken fibroblast cells were transfected with RCAS constructs expressing Luciferase, PDGFB, or Cre. The virus-producing DF1 cells were injected into the brainstem of neonatal mice. Mice were subjected to biweekly bioluminescence imaging to confirm the presence of a brainstem glioma. Upon tumor detection, mice were stratified to various treatment cohorts. (**B**) Schematic showing RCAS-Cre mediated recombination of floxed alleles of *Atm* and tumor suppressor genes (TSG) in *Nestin-expressing* neural progenitor cells that harbor the TVA receptor. Triangles represent loxP sites. (**C**) Axial and (**D**) saggital plane views of the glioma enhanced with gadolinium contrast by MRI. Representative glioma stained with (**E**) hematoxylin and eosin or (**F**) an antibody recognizing the HA tag on PDGFB. All scale bars, 100 μM.

Injection of mice with chicken fibroblasts expressing RCAS-Luciferase enabled tumor development to be confirmed *in vivo* via bioluminescence (Figure 1A). To verify that bioluminescence in the brain was indicative of a tumor, we performed magnetic resonance imaging (MRI) on a subset of mice with detectable luciferase activity. Mice with bioluminescence signal in the brain 4.5 weeks post injection had well-defined tumors within the brainstem visualized by MRI (Figure 1C-D). The tumors enhanced after intravenous injection of gadolinium contrast, indicating that the blood-brain-barrier was compromised (Figure 1C-D). Furthermore, the presence of primary gliomas overexpressing PDGFB was confirmed by immunohistochemistry for the hemagglutinin (HA) tag on PDGFB (Figure 1E-F).

### Deletion of Atm radiosensitizes p53 deficient brainstem gliomas

Exome sequencing of DIPG samples from patients have revealed that mutations in the tumor suppressor *p53* occur in the majority of tumors (21, 22). Likewise, molecular alterations in the *PDGF receptor alpha (PDGFRA)* are well documented (23–25). To model PDGF-driven, *p53* deficient brainstem gliomas, we initiated tumor development in *Nestin^TVA^; p53^FL/FL^; **Atm^FL/+^*** (*nPA^FL/+^*) and *Nestin^TVA^; p53^FL/FL^; **Atm^FL/FL^*** (*nPA^FL/FL^*) mice. *nPA^FL/+^* mice retained expression of one wild-type allele of *Atm* after Cre-mediated recombination and served as the control for all experiments, while *nPA^FL/FL^* mice lacked ATM function. Cre-mediated recombination of floxed alleles of *p53* and *Atm* were confirmed by performing droplet digital PCR on genomic DNA from primary glioma cells sorted by flow cytometry (Supplemental Figure 1 and Figure 2A).

**Figure 2.**
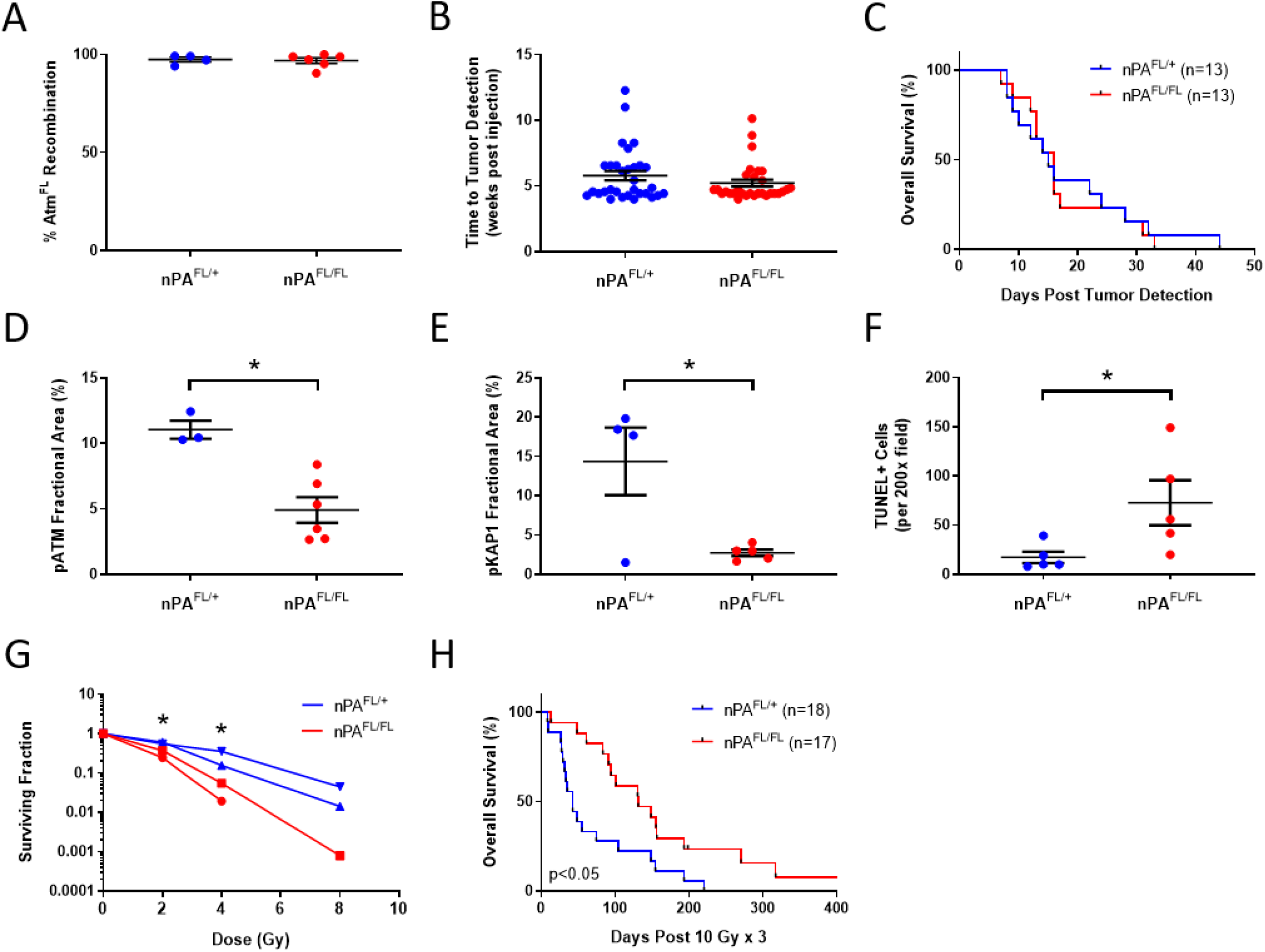
Deletion of *Atm* in *p53* Deficient Gliomas Improves Tumor Response to Radiation. (**A**) Percentage recombination of the floxed allele of *Atm* in flow-sorted glioma cells as assessed by droplet digital PCR. (**B**) Time to tumor development post injection in *nPA^FL/+^* and *nPA^FL/FL^* mice, as detected through *in vivo* bioluminescence imaging. Each circle represents one mouse. (**C**) Kaplan-Meier plot of overall survival in tumor-bearing mice post tumor detection. Quantification of the fractional area of positive staining for (**D**) pATM 2 hours post 10 Gy or (**E**) pKAP1 1 hour post 10 Gy in primary gliomas. (**F**) Number of TUNEL-positive cells in *nPA^FL/+^* and *nPA^FL/FL^* tumors 24 hours post 10 Gy whole brain irradiation. (**G**) Clonogenic survival assay using stroma-depleted *nPA^FL/+^* and *nPA^FL/FL^* tumor cell lines (n=2 cell lines per genotype). (**H**) Kaplan-Meier plot of overall survival in tumor-bearing mice following 3 daily fractions of 10 Gy radiation therapy delivered to the whole brain. \**P*<0.05.

A recent study reported that knocking out *ATM* in tumor cells decreased colony formation in soft agar and slowed tumor growth in immunocompromised mice (26). In a primary brainstem glioma model, however, we detected no difference in the time to tumor detection by *in vivo* bioluminescence imaging between the *Atm* expressing and *Atm* deficient gliomas (Figure 2B). Likewise, deletion of *Atm* in the glioma cells did not impact overall survival of tumor-bearing mice in the absence of radiotherapy (Figure 2C).

To characterize ATM signaling in these brainstem gliomas, we performed immunohistochemistry after radiation therapy. The number of cells staining positively for the activated and phosphorylated form of ATM (pATM) or the ATM target, KRAB-Associated Protein-1 (pKAP1), was significantly diminished post 10 Gy in *nPA^FL/FL^* gliomas when compared to *nPA^FL/+^* gliomas (Figure 2D-E and Supplemental Figure 2). These data provide independent validation of *Atm* recombination in brainstem glioma cells *in vivo,* which resulted in disruption of ATM signaling after radiation therapy.

p53 deficient glioma cells lacking both *Atm* alleles displayed heightened radiosensitivity *in vivo* and *in vitro.* The disruption of ATM signaling resulted in enhanced tumor cell death 24 hours post 10 Gy whole brain irradiation, as detected by TUNEL staining (Figure 2F and Supplemental Figure 2). Similarly, early passage cell lines from tumors in *nPA^FL/FL^* mice were significantly more sensitive to radiation than cell lines derived from tumors in *nPA^FL/+^* mice in clonogenic survival assays *in vitro* (Figure 2G). To investigate whether enhancing tumor cell radiosensitivity improves the survival of mice with primary gliomas treated with radiotherapy, we treated *nPA^FL/+^* and *nPA^FL/FL^* mice with 3 daily fractions of 10 Gy to the brain. The elevated radiosensitivity of glioma cells in *nPA^FL/FL^* mice translated to a tripling of the median survival following radiation therapy (Figure 2H). These data indicate that targeting ATM represents a promising therapeutic approach to improve the response of brainstem gliomas to radiation therapy.

### Deletion of Atm in p53 wild-type gliomas does not improve radiation response

Biddlestone-Thorpe et al. reported that mutations in *p53* were required for radiosensitization by ATM inhibition in a xenograft model of glioblastoma (27). To assess whether loss of p53 signaling was also required for radiosensitization of primary gliomas lacking *Atm* expression, we generated *p53* wild-type tumors by overexpressing *PDGFB* and recombining both alleles of the tumor suppressor *Ink4a/Arf.* Tumor development was initiated in *Nestin^TVA^; Ink4a/Arf^FL/FL^; **Atm^FL/+^*** (*nIA^FL/+^*) and *Nestin^TVA^; Ink4a/Arf^FL/FL^; **Atm^FL/FL^*** (*nIA^FL/FL^*) mice. Although p19^ARF^, an alternate reading frame product from the *Ink4a/Arf* locus, functions to stabilize p53 in response to oncogenic stress, *Ink4a/Arf* deficient tumor cells in this model retain the ability to induce and activate p53 in response to radiation (Supplemental Figure 3). After verifying that the *Ink4a/Arf* deficient cells were functionally p53 intact, we quantified the recombination efficiency of floxed *Atm* and *Ink4a/Arf* alleles using droplet digital PCR. Most flow-sorted primary glioma cells demonstrated full recombination of the floxed *Ink4a/Arf* alleles (Supplemental Figure 4) and there was similar recombination efficiency of *Atm* in the tumors from *nIA^FL/+^* and *nIA^FL/FL^* mice (Figure 3A).

**Figure 3.**
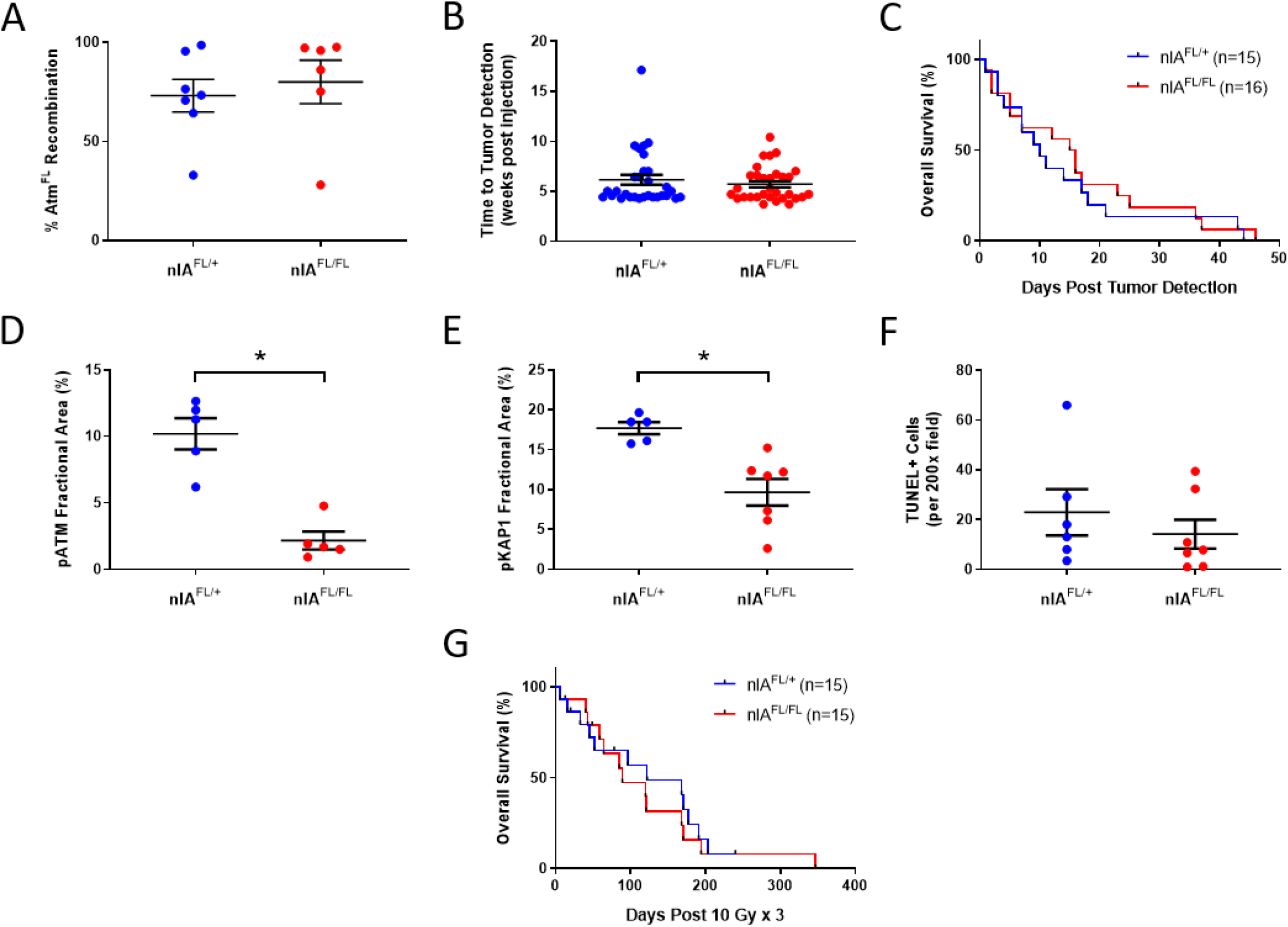
Deletion of *Atm d*oes not Radiosensitize *p53* Wild-Type Gliomas. (**A**) Percentage recombination of the floxed allele of *Atm* in flow-sorted glioma cells as assessed by droplet digital PCR. (**B**) Time to tumor development post injection in *nIA^FL/+^* and *nIA^FL/FL^* mice, as detected through *in vivo* bioluminescence imaging. (**C**) Kaplan-Meier plot of overall survival in tumor-bearing mice post tumor detection in the absence of radiotherapy. Quantification of the fractional area of positive staining for (**D**) pATM 2 hours post 10 Gy or (**E**) pKAP1 1 hour post 10 Gy in primary brainstem gliomas. (**F**) Number of TUNEL-positive cells in *nIA^FL/+^* and *nIA^FL/FL^* tumors 24 hours post 10 Gy whole brain irradiation. (**G**) Kaplan-Meier plot of overall survival in tumor-bearing mice following 3 daily fractions of 10 Gy radiation therapy delivered to the whole brain. \**P*<0.05.

In order to evaluate the impact of *Atm* deletion on growth of *p53* wild-type gliomas, we characterized primary tumor development in *nIA^FL/+^* and *nIA^FL/FL^* mice. Consistent with data generated in the p53-deficient model, deletion of *Atm* did not significantly alter time to tumor detection by *in vivo* imaging (Figure 3B) or the overall survival of tumor-bearing mice in the absence of radiation (Figure 3C).

We also utilized immunohistochemistry to characterize ATM signaling in primary *p53* wild-type tumors post radiation. Staining for pATM and the downstream target pKAP1 was reduced significantly post 10 Gy radiation exposure in *nIA^FL/FL^* gliomas compared to *nIA^FL/+^* gliomas that retained *Atm* expression (Figure 3D-E and Supplemental Figure 5). Despite the significant disruption of ATM signaling following radiation in *nIA^FL/FL^* gliomas, we did not detect an increase in radiation-induced cell death by TUNEL staining (Figure 3F). Furthermore, deletion of *Atm* in *p53* wild-type gliomas did not translate to improved survival following radiotherapy *in vivo* (Figure 3G). Therefore, lack of functional ATM regulates the response of primary brainstem gliomas to radiation therapy in *p53* deficient but not *Ink4a/Arf* deficient (*p53* wild-type) tumors.

### p53 wild-type brainstem gliomas display enhanced radiosensitivity

A recent study using patient-derived DIPG cells and a retrospective cohort of DIPG patients revealed that mutations in *p53* drive radioresistance (5). Most of the patients with *p53* wild-type DIPGs in this study were classified as good clinical and radiological responders after radiation therapy, while only a minority of *p53* mutant tumors could be classified similarly. To determine whether *p53* status influences radiation resistance in the primary models of brainstem glioma, we compared the radiation response of *p53* deficient and wild-type gliomas in *Atm^FL/+^* mice that retained *Atm* expression. While survival of mice with *p53* deficient tumors tripled in response to radiotherapy (27 day increase in median survival), radiotherapy resulted in a twelve-fold increase in the survival of mice bearing *p53* wild-type tumors (110 day increase in median survival) (Figure 4A-B). Importantly, primary gliomas driven by loss of either *p53* or *Ink4a/Arf* grew at a similar rate in the absence of radiation therapy (Figure 4C). However, following three daily fractions of whole brain irradiation mice with *p53* wild-type tumors survived longer than mice with *p53* deficient tumors (Figure 4D). The survival of mice bearing *p53* wild-type gliomas was comparable to the survival of *nPA^FL/FL^* mice with tumors lacking *Atm*. Collectively, these data suggest that *p53* wild-type tumors driven by loss of *Ink4a/Arf* had enhanced sensitivity to radiation therapy (Table 1). These radiosensitive brainstem gliomas could not be further radiosensitized by loss of *Atm*.

**Figure 4.**
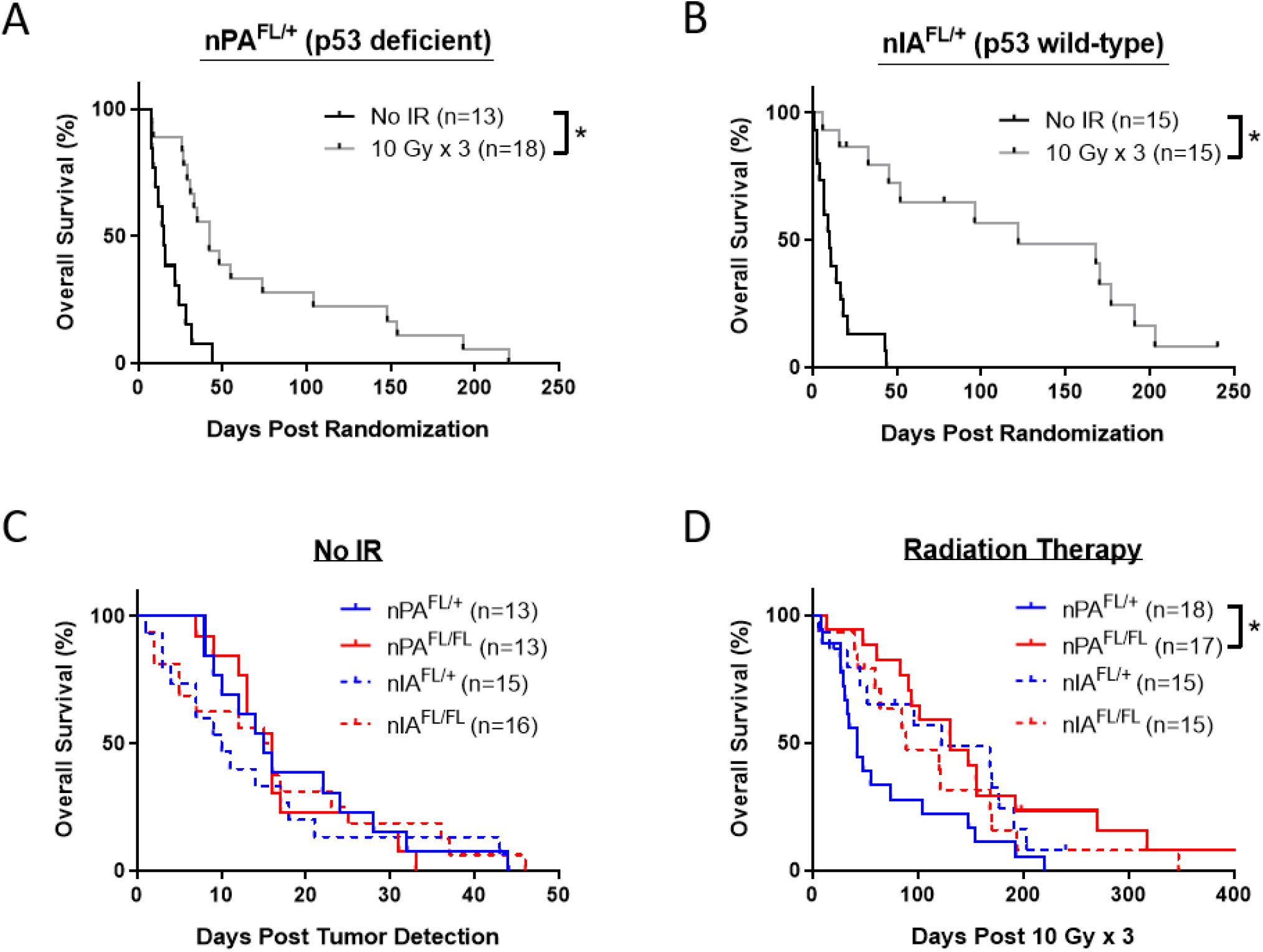
*p53* Wild-Type Gliomas with Intact ATM Function are Sensitive to Radiation. Kaplan-Meier plot of overall survival in (**A**) *p53* deficient tumor-bearing *nPA^FL/+^* mice or (**B**) *p53* wild-type tumor-bearing *nIA^FL/+^* mice that received no radiotherapy or were treated with 3 daily fractions of 10 Gy to the whole brain. (**C**) Kaplan-Meier plot comparing the survival of unirradiated mice with p53 wild-type and deficient brainstem gliomas. (**D**) Kaplan-Meier plot comparing the survival of mice with p53 wild-type and deficient gliomas treated with fractionated radiotherapy. The animals included in these survival curves are the same animals from the survival studies in Figures 2 and 3. \**P*<0.05.

**Table 1.** Radiation Response of Brainstem Glioma GEMMs.

| **Mouse Strain** | **p53** | **INK4a/ARF** | **ATM** | **Radiosensitivity** | **Median Survival (days)** | **Median Survival Post RT (days)** |
| --- | --- | --- | --- | --- | --- | --- |
| **nPA<sup>FL/+</sup>** | - | + | + | Resistant | 15 | 42 |
| **nPA<sup>FL/FL</sup>** | - | + | - | Sensitive | 16 | 131 |
| **nIA<sup>FL/+</sup>** | + | - | + | Sensitive | 10 | 122 |
| **nIA<sup>FL/FL</sup>** | + | - | - | Sensitive | 15.5 | 89 |
| **nPIA<sup>FL/+</sup>** | - | - | + | Very Resistant | 16 | 28 |
| **nPIA<sup>FL/FL</sup>** | - | - | - | Resistant | 21.5 | 44 |

### Proapoptotic genes are induced after radiation in p53 wild-type gliomas

We hypothesized that the radiosensitivity of *Ink4a/Arf* deficient tumors may be caused by the transactivation of p53 targets genes in response to radiation. To evaluate whether genes associated with p53-mediated cell death are induced in primary glioma cells, we quantified the expression of *Bax*, *Puma,* and *Noxa* using real-time PCR. These proapoptotic genes were induced 4 hours post radiation exclusively in the *Ink4a/Arf* deficient cells (Figure 5A-C). In order to confirm that that the radiation-induced expression of these proapoptotic genes was driven by p53, we generated tumors lacking both *p53* and *Ink4a/Arf* in *Nestin^TVA^; p53^FL/FL^; Ink4a/Arf^FL/FL^; **Atm^FL/+^*** (*nPIA^FL/+^*) mice. Deletion of *p53* in the *Ink4a/Arf* deficient gliomas prevented the induction of *Bax*, *Puma,* and *Noxa* after radiation (Figure 5A-C). Similarly, an increase in cleaved caspase-3 following irradiation was observed in *p53* wild-type glioma cells as compared to cells lacking both *p53* and *Ink4a/Arf* (Figure 5D). These data suggest that the radiosensitivity of *p53* wild-type brainstem gliomas may be mediated at least in part by the p53-dependent accumulation of proapoptotic proteins.

**Figure 5.**
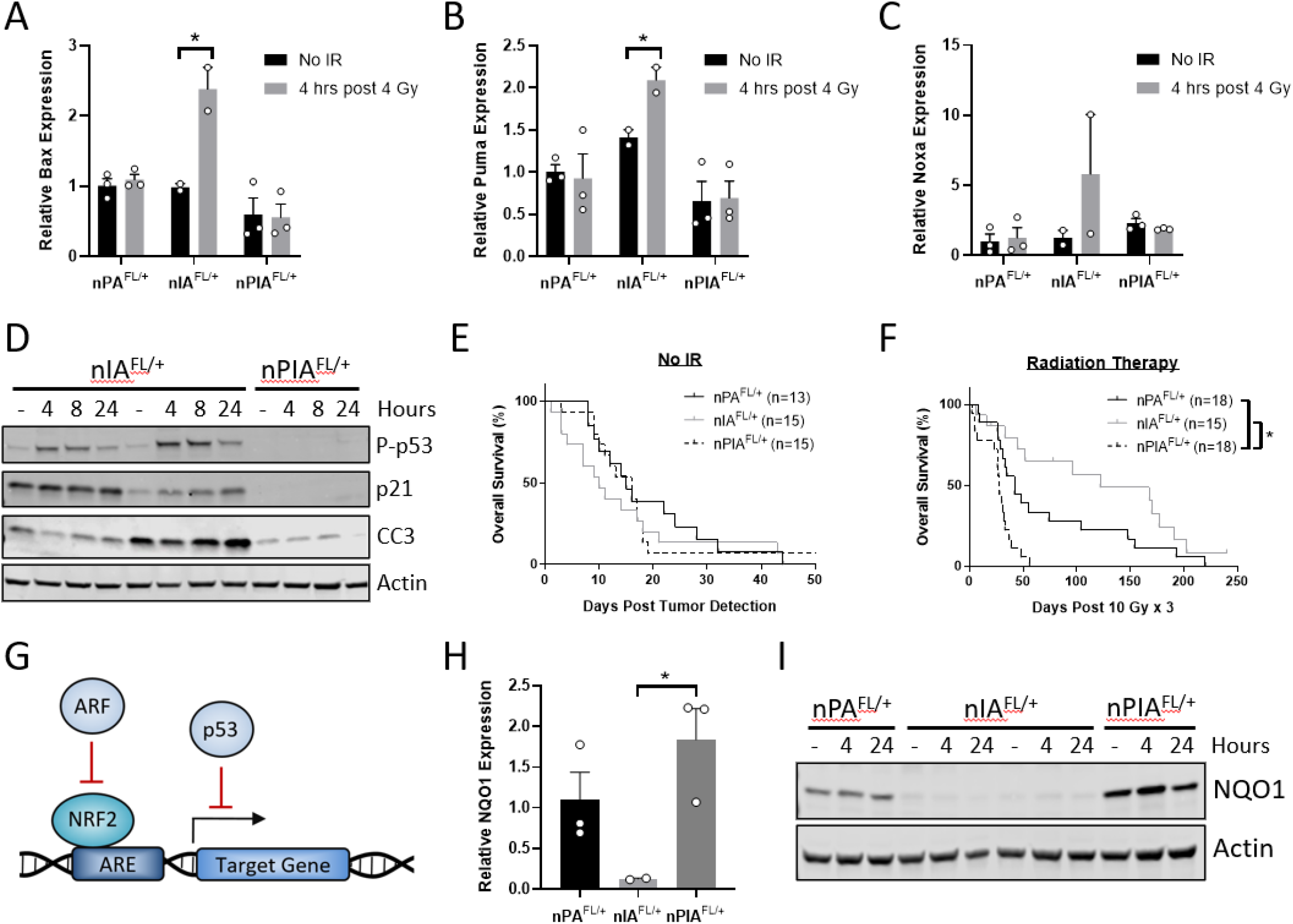
p53 Signaling Mediates Radiosensitivity of *Ink4a/Arf* Deficient Gliomas. Expression of the proapoptotic genes (**A**) *Bax,* (**B**) *Puma,* and (**C**) *Noxa* as quantified by real-time PCR in brainstem glioma cell lines 0 or 4 hours post 4 Gy. (**D**) Western blot showing protein levels of phosphorylated p53 (P-p53), p21, cleaved caspase-3 (CC3), and Actin in *nIA^FL/+^* (n=2 cell lines) and *nPIA^FL/+^* tumor cell lines at various timepoints post 4 Gy. Kaplan-Meier plot comparing the survival of *nPA^FL/+^, nIA^FL/+^,* and *nPIA^FL/+^* mice following (**E**) tumor detection or (**F**) treatment with 3 daily fractions of 10 Gy delivered to the whole brain. Control *nPA^FL/+^* and *nIA^FL/+^* curves were taken from Figures 2 and 3. (**G**) Schematic depicting the negative regulation of the NRF2 signaling pathway by ARF and p53. ARF directly binds to NRF2 and prevents its association with the cognate ARE. p53 mediates transrepression of several NRF2 targets including *Nqo1* by binding to ARE containing promoters. (**H**) Expression of *Nqo1* in primary glioma cell lines as quantified by real-time PCR. (**I**) Western blot showing NQO1 and Actin protein in *nPA^FL/+^, nIA^FL/+^* (n=2 cell lines), and *nPIA^FL/+^* tumor cell lines post treatment with 4 Gy. \**P*<0.05.

### Brainstem gliomas lacking both p53 and Ink4a/Arf are resistant to radiation

To test whether the transactivation of p53 target genes were responsible for the enhanced radiosensitivity of *Ink4a/Arf* deficient brainstem gliomas, we followed tumor-bearing *nPIA^FL/+^* mice for survival with and without radiation exposure. The deletion of *p53* in the *Ink4a/Arf* deficient gliomas did not accelerate tumor growth in the absence of radiation (Figure 5E). However, the disruption of p53 signaling after radiation attenuated the radiosensitivity of *Ink4a/Arf* deficient tumors and significantly reduced the survival of tumor-bearing mice (Figure 5F). Surprisingly, the brainstem gliomas lacking both tumor suppressors were more resistant to radiation than the *p53* deficient gliomas, as indicated by a significant decrease in survival following fractionated radiotherapy (Figure 5F and Table 1). These results suggest that p53 and INK4A/ARF independently contribute to cell survival in response to radiotherapy.

Dysregulation of the NRF2 pathway is an established mechanism of radiation resistance of human tumors and mouse models of cancer (28, 29). NRF2 acts as a master transcriptional regulator of the cellular response to oxidative stress and alters the response to radiation by reducing intracellular reactive oxygen species (ROS) and DNA damage (28, 30). Furthermore, a recent study found that NRF2 and its transcriptional target *SLC7A11* are negatively regulated by p53 and ARF through independent mechanisms (31). While ARF directly interacts with NRF2 and prevents it from binding to the cognate antioxidant response element (ARE), p53 appears to repress transcription of NRF2 targets by binding to ARE containing promoters (32, 33). Therefore, we reasoned that the enhanced radioresistance of brainstem gliomas lacking both *p53* and *Ink4a/Arf* may be mediated by hyperactivation of NRF2 target genes (Figure 5G).

To test this hypothesis, we utilized real-time PCR to examine the expression of the NRF2 target gene *Nqo1.* We found that *p53* deficient glioma cells had elevated expression of *Nqo1* compared to *p53* wild-type cells (Figure 5H). Deletion of *Ink4a/Arf* in addition to *p53* further increased *Nqo1* expression, suggesting that both p53 and INK4A/ARF function to negatively regulate this NRF2 target. We also observed accumulation of NQO1 protein in *p53* and *Ink4a/Arf* deficient glioma cells (Figure 5I). Importantly, the *p53* wild-type tumors lacked *Nqo1* mRNA and protein, correlating with their enhanced radiosensitivity. These data support the hypothesis that hyperactivation of the NRF2 pathway is associated with the increased radioresistance of *nPA^FL/+^* and *nPIA^FL/+^* brainstem gliomas.

### Deletion of Atm reduces the radioresistance of gliomas lacking p53 and Ink4a/Arf

To determine whether targeting ATM can overcome the marked radioresistance of tumors lacking both *p53* and *Ink4a/Arf,* we initiated tumorigenesis in *nPIA^FL/+^* and *Nestin^TVA^; p53^FL/FL^; Ink4a/Arf^FL/FL^; **Atm^FL/FL^** (nPIA^FL/FL^)* mice. Consistent with our previous results, deletion of *Atm* in primary brainstem gliomas did not impact time to tumor formation (Figure 6A) or accelerate tumorigenesis after detection (Figure 6B). We assessed the recombination of floxed alleles of *Atm* by staining tumor tissue for pATM following irradiation with 10 Gy. Activation of ATM was significantly decreased in *nPIA^FL/FL^* tumors compared to *nPIA^FL/+^* tumors where expression of one wild-type allele of *Atm* was retained (Figure 6C and Supplemental Figure 6).

**Figure 6.**
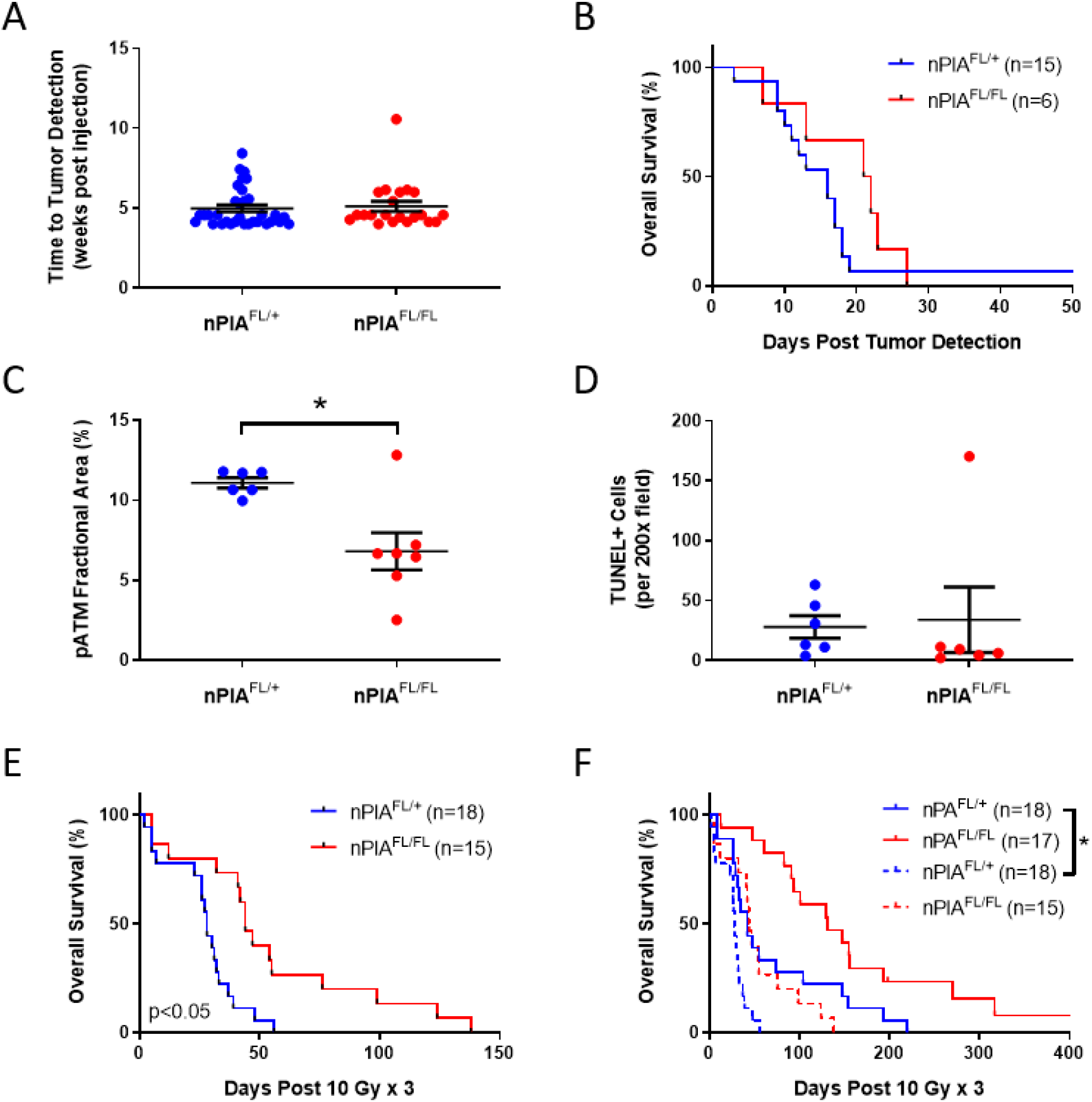
Deletion of *Atm* Improves the Radiation Response of Gliomas Driven by Loss of *p53* and *Ink4a/Arf.* (**A**) Time to tumor development post injection in *nPIA^FL/+^* and *nPIA^FL/FL^* mice, as detected through *in vivo* bioluminescence imaging. (**B**) Kaplan-Meier plot of overall survival in tumor-bearing mice post tumor detection. (**C**) Quantification of the fractional area of positive staining for pATM 2 hours post 10 Gy in primary brainstem gliomas. (**D**) Number of TUNEL-positive cells in *nPIA^FL/+^* and *nPIA^FL/FL^* tumors 24 hours post 10 Gy whole brain irradiation. (**E**) Kaplan-Meier plot of overall survival in tumor-bearing mice following 3 daily fractions of 10 Gy radiation therapy delivered to the whole brain. (**F**) Kaplan-Meier plot comparing the survival of mice with *p53* deficient gliomas and gliomas lacking both *p53* and *Ink4a/Arf* that were treated with fractionated radiotherapy. Control *nPA^FL/+^* and *nPA^FL/FL^* curves were taken from Figure 2. \**P*<0.05.

Despite effective disruption of ATM signaling in *nPIA^FL/FL^* mice, we did not observe an increase in cell death 24 hours after radiation exposure (Figure 6D and Supplemental Figure 6). Nevertheless, the deletion of *Atm* modestly improved the survival of tumor-bearing mice treated with fractionated radiotherapy (Figure 6E). These results further demonstrate that in primary brainstem gliomas, the status of *p53* is a critical determinant of tumor response to concurrent radiotherapy and inactivation of ATM. Interestingly, the survival after irradiation of *nPIA^FL/FL^* mice with tumors lacking both copies of *p53*, *Ink4a/Arf* and *Atm* was comparable to *nPA^FL/+^* mice whose tumors lacked *p53* but retained expression of *Ink4a/Arf* and *Atm* (Table 1), indicating that *Atm* deficient gliomas driven by loss of *p53* and *Ink4a/Arf* remain relatively radioresistant (Figure 6F). Whereas deletion of *Atm* tripled the radiation response of *p53* deficient tumors that retained *Ink4a/Arf* expression, the same genetic modification only increased survival by 1.5-fold in tumors lacking both *p53* and *Ink4a/Arf* (Table 1). Therefore, other genetic alterations can limit the ability of ATM inactivation to fully radiosensitize *p53* deficient gliomas.

## Discussion

To efficiently characterize and modulate the radioresistance of DIPGs, we generated GEMMs of brainstem glioma by combining Cre-*loxP* and RCAS-TVA technology (20). These sophisticated techniques enabled precise control over the cell-of-origin and the genetic modifications driving gliomagenesis (34). As genetic alterations in *PDGFRA* and the tumor suppressor *p53* are two of the most common genetic alterations in human DIPGs, we first investigated radiation response in a primary brainstem glioma model overexpressing *PDGFB* and lacking *p53.* We also generated a second model in which p53 function was retained but *Ink4a/Arf* was deleted in *Nestin-expressing* progenitor cells. Although mutations in the *CDKN2A* locus are rare in DIPGs, this locus is epigenetically repressed by the prevalent K27M mutation in histone H3.3 (35). In future studies, we will assess the impact of the H3.3K27M mutation on radiation response of DIPGs as well as the response to *Atm* deletion. However, it is noteworthy that a recent shRNA screen identified ATM as a promising target for the radiosensitization of H3.3K27M mutated DIPG cells (5).

In addition to incorporating the genetic alterations found in many DIPG patients, primary models of brainstem glioma have been shown to resemble human tumors histologically and to undergo a similar pattern of invasion (36). Furthermore, by delivering fractionated radiotherapy alone to these mouse models, we were able to achieve a modest improvement in survival but not tumor cure, which mimics the clinical response of DIPG patients treated with radiotherapy. Using the various GEMMs, we also showed that *p53* wild-type gliomas were more radiosensitive than *p53* deficient gliomas, in agreement with a retrospective analysis of DIPG patients treated with radiotherapy (5). As our primary tumor models closely phenocopy human brainstem gliomas, these murine models represent valuable tools for identifying drug targets that may improve clinical outcomes in DIPG patients treated with radiation therapy.

Utilizing complimentary *p53* deficient and *p53* wild-type (*Ink4a/Arf* deficient) models of brainstem glioma, we found that disruption of ATM signaling preferentially radiosensitized tumors lacking *p53.* Although other groups have reported that *p53* status regulates radiosensitization by ATM inhibition (27, 37), our study is the first to implicate p53 status in radiosensitization by *Atm* deletion in a primary tumor system. Furthermore, we previously reported that endothelial-cell specific deletion of *Atm* is not sufficient for radiosensitization following focal heart irradiation, but simultaneous loss of *p53* and *Atm* facilitates radiation-induced heart injury (38). Collectively, this body of work suggests that in tumors at specific anatomic sites, targeting ATM my provide a therapeutic window in cancers that have lost p53 function.

As most DIPGs contain *p53* mutations (21, 22), we hypothesize that treatment with ATM inhibitors will selectively radiosensitize *p53* mutant tumor cells without impacting the radiosensitivity of adjacent *p53* wild-type stromal cells and normal tissue. Consistent with this hypothesis, studies in *Atm* knockout mice have shown that deletion of both copies of *Atm* radiosensitizes the intestines and bone marrow, but does not radiosensitize other normal tissues including the brain, skin, lung and heart (16, 39, 40). Further supporting the notion of a therapeutic window for treating children with DIPG with concurrent radiotherapy and ATM inhibition are findings that *Atm* null mice are resistant to radiation-induced cell death in many regions of the central nervous system, including the dentate gyrus, cerebellum, and cerebral cortex (41). Taken together, these results suggest that ATM inhibitors could simultaneously radiosensitize a *p53* mutant tumor in the brainstem while protecting the surrounding normal brain tissue. As ATM inhibitors that penetrate the blood-brain-barrier have been recently developed (19) and are now in clinical trials with concurrent radiotherapy for adults with glioma and brain metastases (NCT03423628), our findings have important implications for the clinical translation of concurrent ATM inhibition and radiotherapy to children with DIPGs.

Our study showed that *p53* wild-type brainstem gliomas respond well to radiotherapy, although they are not further radiosensitized by deletion of *Atm*. The radiosensitivity of these tumors was mediated by p53, as deletion of *p53* completely attenuated the radiation response. Surprisingly, tumors lacking both *p53* and *Ink4a/Arf* were more resistant to radiation than tumors driven by loss of *p53* alone. These findings suggest that *p53* and *Ink4a/Arf* independently regulate tumor response to radiation. Consistent with a recent report highlighting the role of these two tumor suppressors in the negative regulation of the NRF2 pathway (31), we found that expression of *Nqo1* was significantly increased in tumor cells lacking *p53* and *Ink4a/Arf.* These results support a role for *Ink4a/Arf* in tumor development outside of its canonical activity as a regulator of p53 protein stability. In addition to providing a possible mechanism for the heightened radioresistance of *nPIA^FL/+^* gliomas, increased expression of *Nqo1* may also provide a selective pressure in human tumors that acquire mutations in both the *p53* and *CDKN2A* loci (42–45). Interestingly, others have recently reported that substrates for NQO1 that lead to a futile redox cycle using NAD(P)H are effective radiosensitizers (46). Targeting NQO1 during radiotherapy may therefore represent a promising approach to improve outcomes in DIPG in tumors with genotypes that limit radiosensitization by targeting ATM.

## Methods

### Mouse strains

All of the mouse strains used in this study have been described previously, including *Nestin^TVA^, p53^FL^, ATM^FL^,* and *Ink4a/Arf^FL^* mice *(47–50). Nestin^TVA^* mice were provided by Oren Becher (Northwestern University), *p53^FL^* mice were provided by Anton Berns (Netherlands Cancer Institute), *ATM^FL^* mice were provided by Frederick Alt (Boston Children’s Hospital), and *Ink4a/Arf^FL^* mice were provided by Ronald DePinho (MD Anderson Cancer Center). All mouse strains were maintained on a mixed genetic background. Male and female littermate controls were utilized in all experiments to minimize the effect of differences in sex, genetic background, and environment on the experimental endpoint.

### Primary glioma induction

DF1 chicken fibroblast cells (ATCC) were cultured at 39°C in DMEM with high glucose (ATCC) supplemented with 10% fetal bovine serum, 2mM L-glutamine, and 1% antibiotic-antimycotic. The DF1s were transfected with various RCAS plasmids using X-TremeGENE 9 (Roche). Primary brainstem gliomagenesis was induced by injecting *Nestin^TVA^* mice (postnatal days 3-5) with 1×10^5^ virus producing DF1 cells resuspended in 1 μL. A Hamilton syringe was used to inject the DF1 cells into the brainstem as previously described (36). DF1 cells expressing RCAS-PDGFB, RCAS-Cre, and RCAS-Luciferase were injected in equal ratios for all studies except for the fluorescence-activated cell sorting (FACS) experiments. In the FACS experiments, mice were injected with 1×10^5^ virus producing DF1 cells expressing RCAS-PDGFB, RCAS-Cre, RCAS-Luciferase, and RCAS-GFP at a 2:2:1:1 ratio.

### In vivo imaging of primary gliomas

Following chemical hair removal, mice were injected intraperitoneally with 150 mg/kg D-luciferin (Gold Biotechnology). Mice were anesthetized in a chamber with 2-3% isoflurane in oxygen and bioluminescence was assessed 10 minutes following injection of D-luciferin. Mice were monitored biweekly beginning 4.5 weeks post tumor initiation using the IVIS Lumina III (PerkinElmer).

Magnetic resonance imaging (MRI) was performed on a subset of mice using a 7 Tesla preclinical scanner (Bruker BioSpec 7/20, Bruker, Ettlingen, Germany) equipped with cryogenic receivers (Cryoprobe, Bruker). Mice were anaesthetized with isoflurane and the brain was imaged three-dimensionally at 0.15mm isotropic resolution using a *Γ*2-weighted acquisition, with parameters fg-gr and a. *T*1-weighted acquisition. Imaging took place prior to and following injection with a gadolinium-based contrast agent (Magnevist, DOSE) through a tail-vein catheter. The location and size of the brainstem gliomas were determined by manual segmentation.

### Radiation treatment

All primary tumor irradiations were performed with the X-RAD 225Cx small animal image-guided irradiator (Precision X-Ray). Mice were placed in a prone position and anaesthetized with isoflurane. Fluoroscopy with 40 kVp, 2.5 mA X-rays filtered through 2 mm aluminum was utilized to center the radiation field on the target. Mice were irradiated with 225 kVp, 13 mA X-rays filtered through 0.3 mm copper using a 15 x 40 mm rectangular radiation field with an average dose rate of ~280 cGy/min. Tumor bearing mice received whole brain irradiation with right and left lateral fields.

Primary tumor cell lines were irradiated *in vitro* using either the X-RAD 320 or the X-RAD 160 Biological Irradiator (Precision X-ray Inc.). In the X-RAD 320, cells were treated 50 cm from the radiation source with 320 kVP, 10 mA X-rays, using a 2 mm aluminum filter. In the X-RAD 160, cells were treated 40 cm from the radiation source with 160 kVP, 18 mA X-rays, using a 2 mm aluminum filter. Dose rates were calculated by the Radiation Safety Division at Duke University.

### Histological analysis

Hematoxylin and eosin (H&E) staining and immunohistochemistry were performed on both frozen and paraffin-embedded sections. Frozen tissues were first fixed in 4% paraformaldehyde for 24 hours, then transferred to a 30% sucrose solution for 24-48 hours, and finally snap frozen in OCT compound (Sakura Finetek) using a dry ice and isopentane slurry. A cryostat was used to generate 10 μm tissue sections. Paraffin-embedded tissues were fixed in 10% neutralized formalin overnight and preserved in 70% ethanol until embedding in paraffin. Prior to staining, 5 μm tissue sections were deparaffinized in xylene and rehydrated in a graded series of ethanol and water washes.

For immunohistochemistry, endogenous peroxidase activity was blocked by treating sections with 3% H_2_O_2_ and antigens were retrieved with Antigen Unmasking Solution (Vector Laboratories). Tissues were blocked in 10% serum with 0.25% Tween 20 (Sigma Aldrich) prior to incubation in primary antibody overnight. The following primary antibodies were used: rabbit polyclonal to HA-probe (1:250, Santa Cruz, sc-805), mouse monoclonal to Ser1981 phosphorylated ATM (1:200, Millipore Sigma, 05-740), and rabbit anti-mouse Ser824 phosphorylated Kap1 (1:200, Bethyl Laboratories, #A300-767A). The tissues were incubated in biotinylated secondary antibodies for 30 minutes at room temperature and treated with VECTASTAIN Elite ABC Reagent (Vector Laboratories) per the manufacturer’s instructions. The DAB Peroxidase Substrate Kit (Vector Laboratories) was utilized to visualize positive staining prior to counterstaining with Mayer’s hematoxylin and dehydrating in a graded series of ethanol and water. Representative images were acquired with a Leica DFC450 brightfield microscope using Leica Suite software (Leica Microsystems). ImageJ (NIH) was utilized to quantify positively stained cells or fractional areas by a single observer blinded to genotype and treatment.

### Immunofluorescence

Immunofluorescence was performed on both frozen and paraffin-embedded tissue sections, which were prepared as described in the histological analysis section. To perform TUNEL staining, the In Situ Cell Death Detection Kit (Roche) was used per the manufacturer’s instructions. Nuclear labeling was performed with VECTASHIELD mounting medium with DAPI (Vector). Representative images were acquired with a DFC340 FX fluorescence microscope using Leica Suite software (Leica Microsystems). ImageJ (NIH) was utilized to quantify positively stained cells or fractional areas by a single observer blinded to genotype and treatment.

### Isolation of primary glioma cell lines

Primary brainstem gliomas were digested at 37°C for 15 minutes in Dissociation Solution containing Earle’s Balanced Salt Solution (EBSS), 0.94 mg/mL papain (Worthington), 0.18 mg/mL EDTA, 0.18 mg/mL cysteine, and 0.06 mg/mL deoxyribonuclease I (DNase). Following extensive trituration, the glioma cells were pelleted and resuspended in an Ovomucoid Solution, containing 0.7 mg/mL ovomucoid (Worthington) and 0.01 mg/mL DNase in NeuroCult basal medium (StemCell). The glioma cells were cultured in DMEM with high glucose and pyruvate (Gibco) supplemented with 10% fetal bovine serum, 2mM L-glutamine, and 1% antibiotic-antimycotic (Gibco). Cells were passaged at least five times to enrich for tumor cells and deplete stromal cells.

### Clonogenic survival assays

Tumor cells were plated in triplicate at two different cell densities and allowed to adhere overnight. After irradiation, cells were incubated 1-2 weeks until colonies were detected in the unirradiated controls. The cells were washed with PBS, fixed with 70% ethanol, stained with Coomassie Brilliant Blue (Bio-Rad), and rinsed with deionized water. Colonies, defined as collections of at least 50 adjacent cells, were quantified by a single observer to calculate surviving fractions relative to unirradiated controls.

### Fluorescence-activated cell sorting

At the time of tumor initiation, mice were injected with DF1 cells expressing RCAS-GFP as described in the primary glioma induction section. Upon detection of a tumor by bioluminescence imaging, primary glioma cells were isolated and red blood cells were lysed using ammonium-chloride-potassium (ACK) lysing buffer (Lonza). Tumor cells were resuspended in flow buffer (Hank’s Balanced Salt Solution with Ca2+ and Mg2+, 5% fetal bovine serum, 2 mM EDTA) and stained with Zombie Aqua Fixable Viability Kit (BioLegend, 1:200). Living, GFP-positive glioma cells were flow sorted using the Astrios Cell Sorter (Beckman Coulter).

### Droplet digital PCR to quantify recombination

Genomic DNA was extracted from flow sorted primary glioma cells using PicoPure DNA extraction kit (Applied Biosystems) and quantified using NanoDrop (Thermo Scientific). To assess the frequency of Cre-mediated recombination of floxed alleles in the glioma models, we designed hexachloro-fluorescein (HEX) and fluorescein amidite (FAM) conjugated Taqman probes to specifically detect the unrecombined and recombined alleles of *Atm^FL^, p53^FL^,* or *Ink4a/Arf^FL^*. The sequences of primers and probes for these assays were:

*Atm* Recombined Primer: 5’-TCACAACCATCTTCAACCCC-3’

*Atm* Floxed Forward Primer: 5’-AATCATCCTTTAATGTGCCTCC-3’

*Atm* Floxed Reverse Primer: 5’-TTCATCATCGTCGACCGC-3’

*Atm* Recombined Probe: 5’-ACACATGCATGCAGGCAGAGCATCCCT-3’

*Atm* Floxed Probe: 5’-AGCTGTTACTTTTGCGTTTGGTGTGGCG-3’

*p53* Recombined Primer: 5’-TCACCATCACCATGAGACAGG-3’

*p53* Floxed Forward Primer: 5’-TGCCCTCCGTCCTTTTTCG-3’

*p53* Floxed Reverse Primer: 5’-GGACAGCCAGGACTACACAG-3’

*p53* Recombined Probe: 5’-CTTGATATCGAATTCCTGCAGCCCGGG-3’

*p53* Floxed Probe: 5’-ATGCTATACGAAGTTATCTGCAGCCCGG-3’

*Ink4a/Arf* Recombined Primer: 5’-CCTAGAGGTTGATGACAAGG-3’

*Ink4a/Arf* Floxed Forward Primer: 5’-CTGTGGCAGGATATAACTTCG-3’

*Ink4a/Arf* Floxed Reverse Primer: 5’-GAATCCCCATCCACTCTGGA-3’

*Ink4a/Arf* Recombined Probe: 5’-CATTATACGAAGTTATGGCGCGCCC-3’

*Ink4a/Arf* Floxed Probe: 5’-CTCTGAAAACCTCCAGCGTATTCTGGTA-3’

Droplet digital PCR on genomic DNA diluted to 25 ng/uL was performed using the QX200 Droplet Digital PCR System (Bio-Rad) according to the manufacturer’s instructions. Suggested PCR cycling conditions were used: 94°C for 30 seconds and 60°C for 1 min (40 cycles), followed by 98°C for 10 min. Restriction digestion was performed in the droplet digital PCR reaction using HindIII for the *Atm* and *Ink4a/Arf* reactions and CviQI for the *p53* reaction. Data were analyzed using the QuantaSoft software (Bio-Rad) by an investigator blinded to genotype. Only samples primarily containing tumor cells (as defined by full recombination of floxed tumor suppressor alleles) were assessed for recombination of the floxed allele of Atm.

### Immunoblotting

Primary tumor cells were lysed in cold RIPA buffer (Sigma Aldrich) supplemented with Aprotinin (Sigma Aldrich), PhosStop tablet (Sigma Aldrich), cOmplete mini protease inhibitor cocktail tablet (Sigma Aldrich), and 1mM PMSF. Soluble proteins were quantified using the Pierce BCA Protein Assay (ThermoFisher) and equal amounts of protein were resolved by SDS-PAGE. Proteins were transferred to nitrocellulose membranes and immunoblotted with specific antibodies for p19^ARF^ (Novus Biologicals, NB200-174, 1:250), p53 (Cell Signaling Technology, #2524, 1:250), Ser15 phosphorylated p53 (Cell Signaling Technology, #9284, 1:250), p21 (Santa Cruz, sc-471, 1:250), Cleaved Caspase-3 (Cell Signaling, 9661S, 1:1,000), NQO1 (Abcam, ab34173, 1:1,000), Actin (BD Transduction, 612656, 1:10,000) or GAPDH (ProteinTech, 60004-1-IgG, 1:1,000). Proteins were visualized using infrared fluorophore labeled secondary antibodies (Li-Cor Biosciences, IRDye800 and IRDye680) and imaged with the Odyssey imaging system (Li-Cor Biosciences).

### Quantitative real-time PCR

Total RNA was extracted from tumor cells using the Direct-zol RNA Kit (Zymo) and reverse transcription was performed using the iScript Advanced cDNA Synthesis Kit (Bio-Rad). Quantitative real-time PCR was utilized to detect mRNA expression using TaqMan Fast Advanced Master Mix (Thermo Fisher) and TaqMan probes (Thermo Fisher, Mm00451763_m1 for *Noxa,* Mm00432051_m1 for *Bax,* Mm00519268_m1 for *Puma,* Mm01253561_m1 for *NQO1,* Mm99999915_g1 for *Gapdh). Gapdh* expression was used as an internal control for RNA concentration across samples. Every sample was run in triplicate and the results were averaged for each assay.

### Statistics

All data are presented as means ± s.e.m. Student’s *t* test (two-tailed) was utilized to compare the mean of two groups. Two-way ANOVA was utilized to examine the interaction between genotype and radiation or time point followed by Bonferroni’s multiple comparisons test for pairwise comparisons. Kaplan-Meier analysis was performed for the survival studies, followed by the log-rank test for statistical significance. A *P* value less than 0.05 indicated significance. Prism 7 (GraphPad Software, Inc.) was used for the statistical analysis.

### Study Approval

All animal studies were approved by the Institutional Animal Care and Use Committee (IACUC) at Duke University.

## Author Contributions

KMD, OJB, and DGK conceived the study. KMD, MC, OJB, and DGK designed experiments. KMD, BFS, JSM, JB, and MC generated the data. DMC provided training in generating primary brainstem gliomas. KMD and DGK analyzed the data. NTW irradiated the mice. LL tattooed and genotyped the mice. YM performed the histology. KMD and DGK wrote the manuscript. KMD, MC, OJB, and DGK reviewed the manuscript.

## Acknowledgements

This work is dedicated to the memory of Rose Sugarman, who died from a DIPG. We thank John Nouls from the Duke Center for In Vivo Microscopy for assistance in obtaining the MRI. This work was supported by grants to DGK from National Cancer Institute (R35 CA197616), Pediatric Brain Tumor Foundation, Hannah’s Heroes St. Baldrick’s Research Grant, and The Leon Levine Foundation. DGK and KMD were supported by the National Cancer Institute (Duke Brain SPORE P50-CA19099). MC was supported by the National Cancer Institute (F30 CA206424). OJB was supported by Maddox’s Warriors, the Fly A Kite Foundation, Cristian Rivera Foundation, and the Rory David Deutsch Foundation.

