## Supplemental Figures for "Tumor genotype dictates radiosensitization after *Atm* deletion in brainstem gliomas"

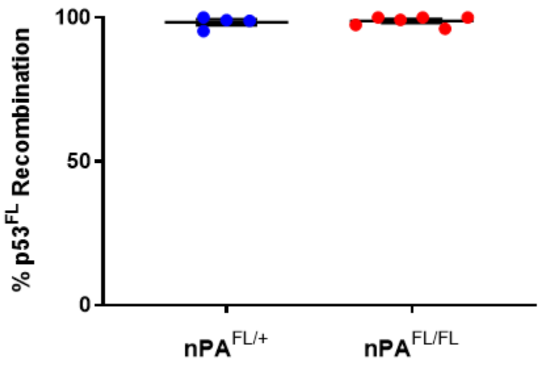


**Supplemental Figure 1. Recombination Efficiency in *p53* Deficient Gliomas.** Percentage recombination of the floxed allele of *p53* in flow-sorted glioma cells as assessed by droplet digital PCR.

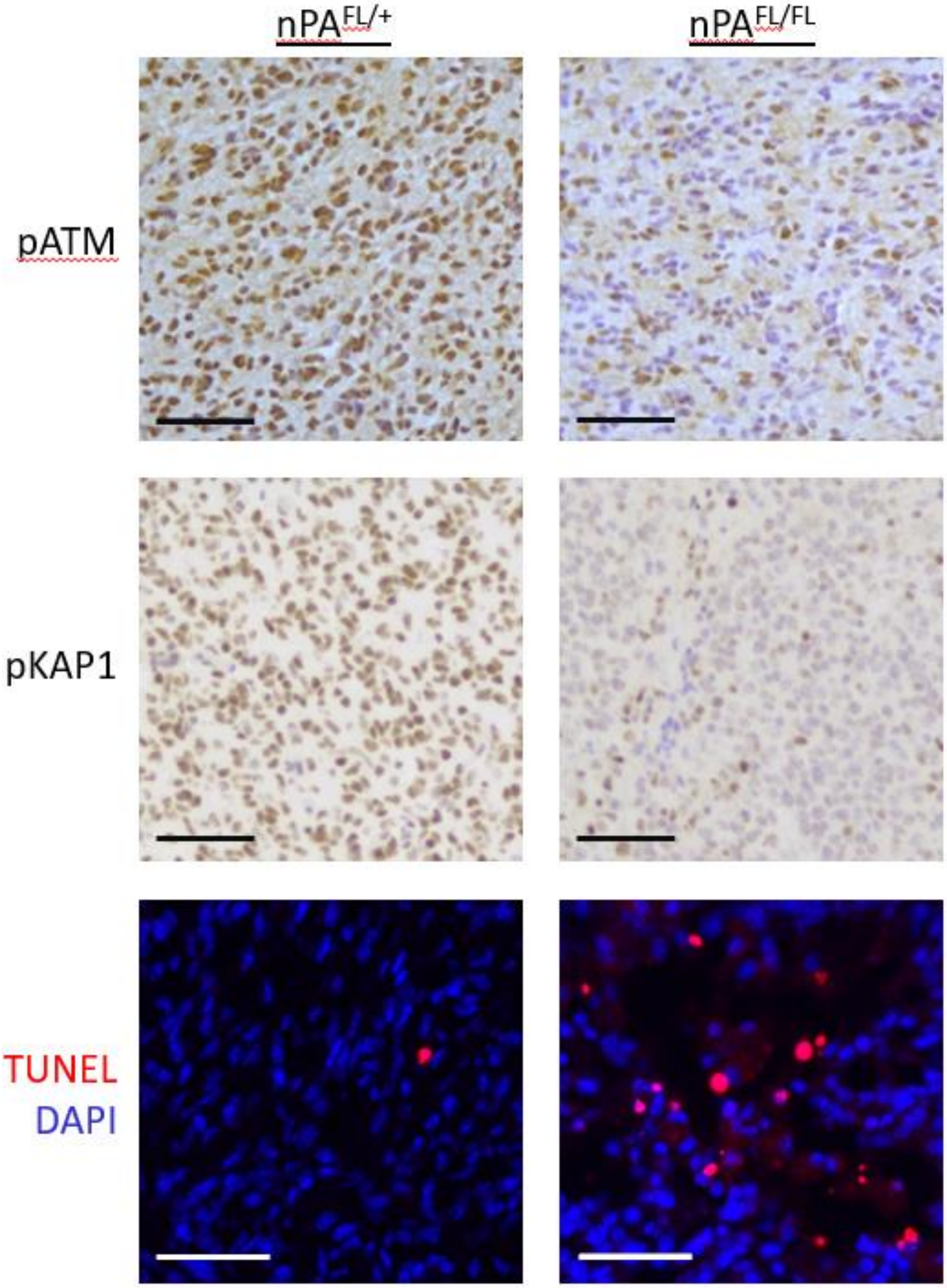


**Supplemental Figure 2. Representative Staining of *p53* Deficient Gliomas.** Representative images of pATM, pKAP1, and TUNEL staining in brainstem gliomas isolated from *nPA*$^{FL/+}$ (left) or *nPA*$^{FL/FL}$ (right) mice. All scale bars, 50 µM.

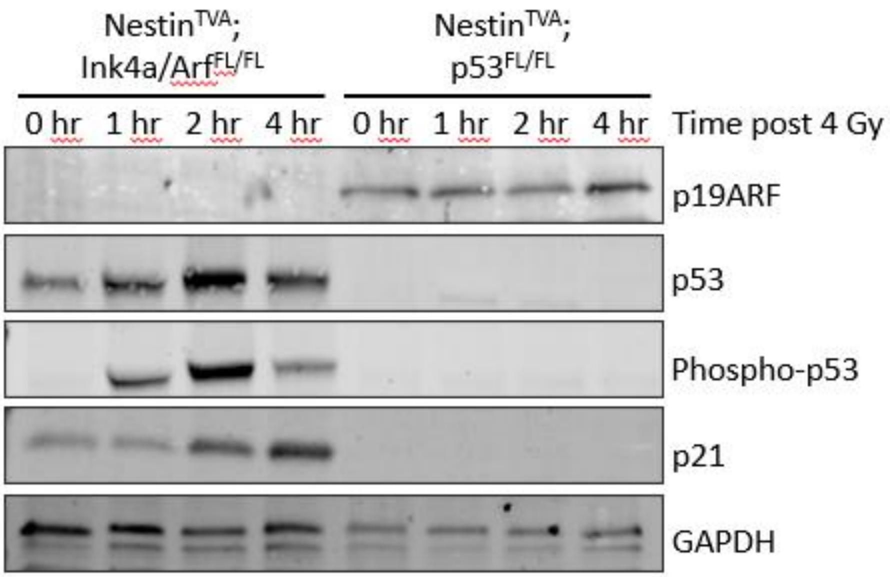


**Supplemental Figure 3. Radiation-Induced p53 Signaling in *Ink4a/Arf* Deficient Gliomas.**

Western blot showing protein levels of $p19^{ARF}$, p53, phosphorylated p53, p21, and GAPDH in *p53* wild-type (from *Ink4a/Arf*$^{FL/FL}$ mice) and *p53* deficient (from *p53*$^{FL/FL}$ mice) primary tumor cell lines at the indicated timepoints post treatment with 4 Gy.

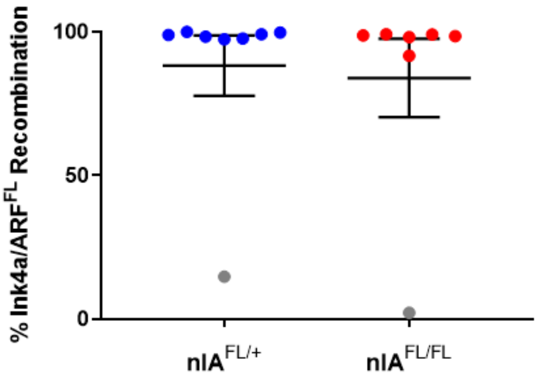


**Supplemental Figure 4. Recombination Efficiency in *p53* Wild-Type Gliomas.** Percentage recombination of the floxed allele of *Ink4a/Arf* in flow-sorted glioma cells as assessed by droplet digital PCR. Grey circles represent samples with poor recombination of *Ink4a/Arf* $^{FL}$ alleles. These samples were excluded from further characterization of the floxed allele of *Atm* by droplet digital PCR.

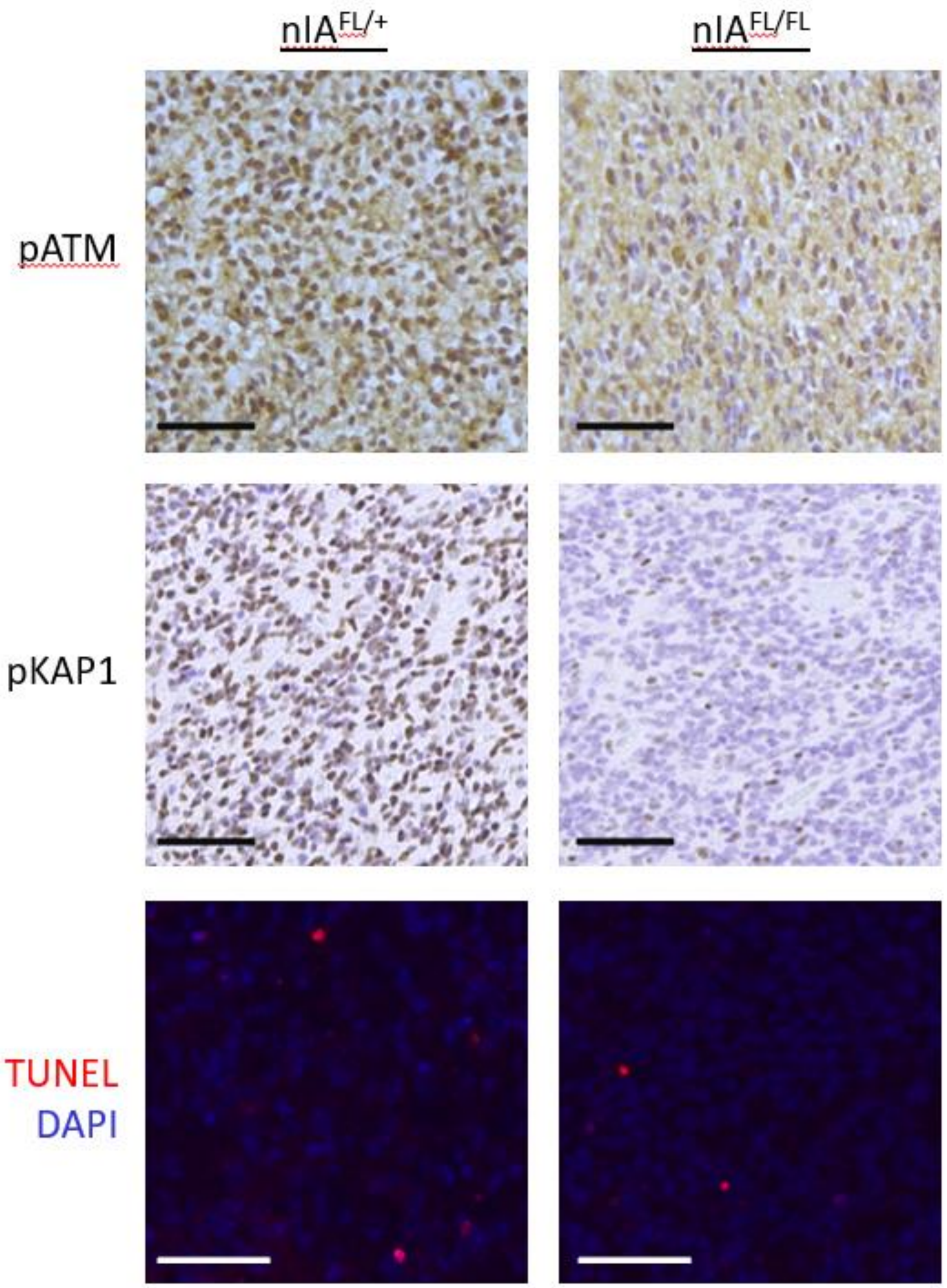


**Supplemental Figure 5. Representative Staining of *p53* Wild-Type Gliomas.** Representative images of pATM, pKAP1, and TUNEL staining in brainstem gliomas isolated from $nIA^{FL/+}$ (left) or $nIA^{FL/FL}$ (right) mice. All scale bars, 50 μM.

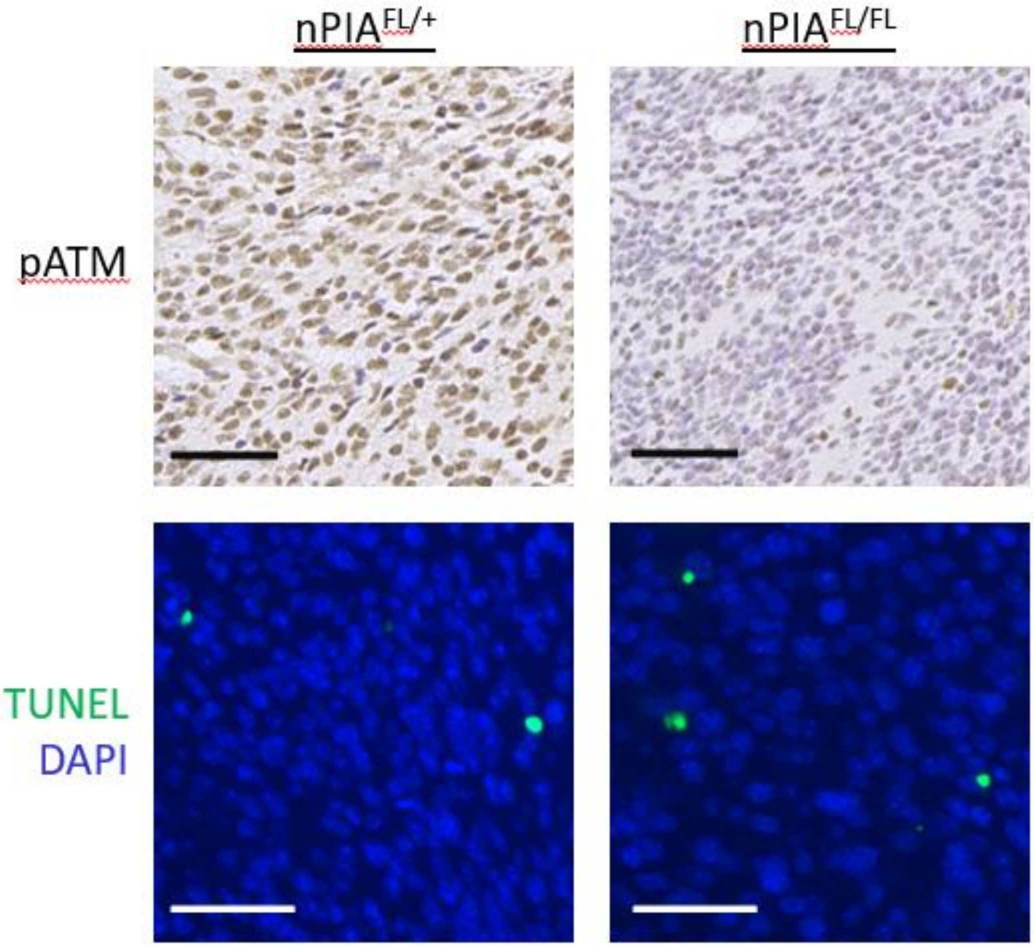


**Supplemental Figure 6. Representative Staining of *p53* and *Ink4a/Arf* Deficient Gliomas.**

Representative images of pATM and TUNEL staining in brainstem gliomas isolated from *nPIA*$^{FL/+}$ (left) or *nPIA*$^{FL/FL}$ (right) mice. All scale bars, 50 µM.
